# Introduction of Surface Loops as a Tool for Encapsulin Functionalization

**DOI:** 10.1101/2021.05.13.444092

**Authors:** Sandra Michel-Souzy, Naomi M. Hamelmann, Sara Zarzuela-Pura, Jos M. J. Paulusse, Jeroen J. L. M. Cornelissen

## Abstract

Encapsulin based protein cages are nanoparticles with different biomedical applications, such as targeted drug delivery or imaging agents. These particles are biocompatible and can be produced in bacteria, allowing large scale production and protein engineering. In order to use these bacterial nanocages in different applications, it is important to further explore the potential of their surface modification and optimize their production. In this study we design and show new surface modifications of the *Thermotoga maritima* (Tm) and *Brevibacterium linens* (Bl) encapsulins. Two new loops on Tm encapsulin with a His-tag insertion after the residue 64 and the residue 127, and the modification of the C-terminal on Bl encapsulin, are reported. The multi-modification of the Tm encapsulin enables up to 240 different functionalities on the cage surface, resulting from 4 potential modifications per protein subunit. We furthermore report an improved protocol giving a better stability and providing a notable increase of the production yield of the cages. Finally, we tested the stability of different encapsulin variants over a year and the results show a difference in stability arising from the tag insertion position. These first insights in the structure-property relationship of encapsulins, with respect to the position of a function loop, allow for further study of the use of these protein nanocages in biomedical applications.

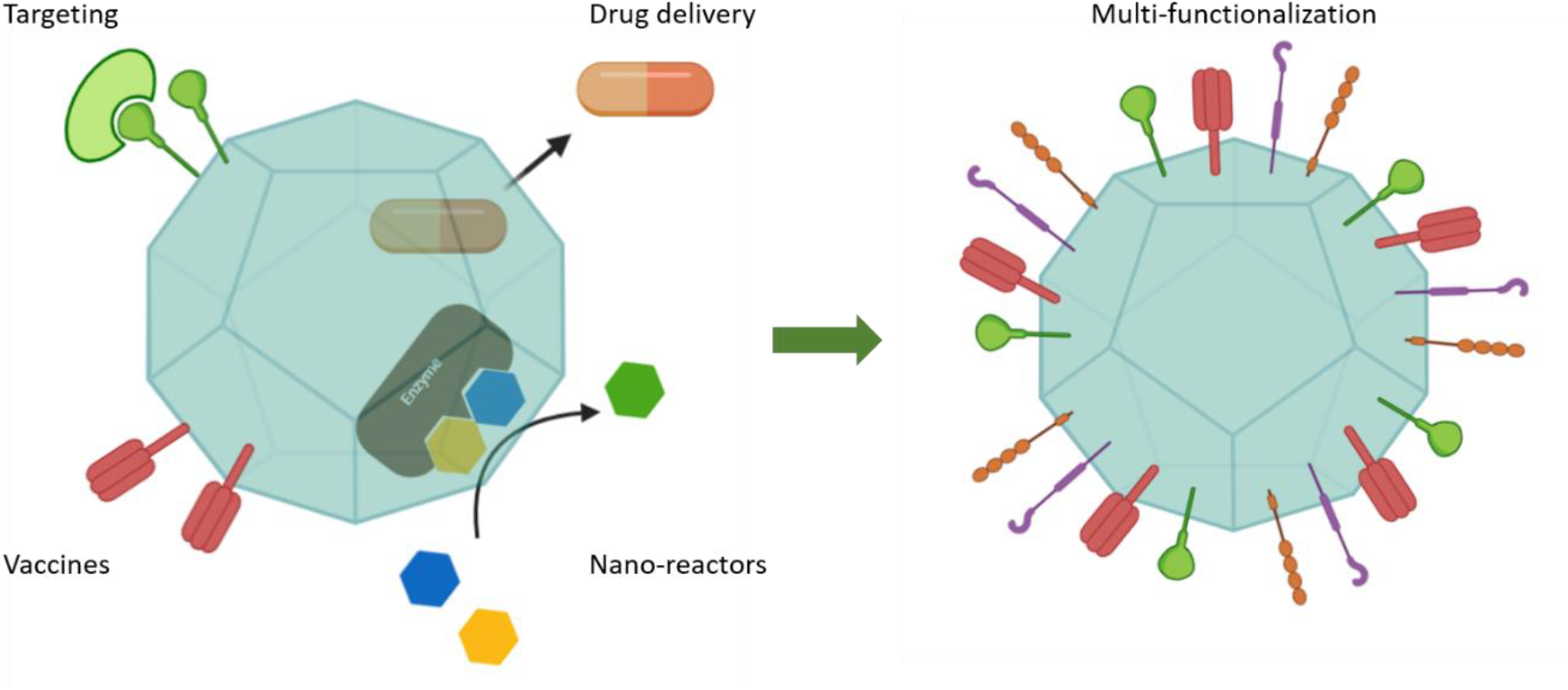

## INTRODUCTION

The constant development of biomedical tools for drug targeting, drug delivery, vaccines or imaging requires more efficient biocompatible carriers. Thus, nanoparticles have been built such as liposomes,^1^ polymers,^2^ micelles,^3^ or protein cages.^4^ Previous studies on protein cages can be found for a variety of different applications such as targeted drug delivery, gene reporting and vaccines.^5–7^ Protein cages are well-defined monodisperse, hollow structures and they are biocompatible thanks to the fact that they are protein based, with a spherical shape, in the range of 1-100nm size.^8–10^ Among the variety of protein cages, our work focuses on encapsulin cages which were discovered in some bacteria and archaea^11^ and may have viral origin shared with the capsid proteins of tailed bacteriophages.^12–14^ The advantages of bacterial protein cages is that they can be easily engineered, are easy to produce and they self-assemble in bacteria.^15^ Furthermore, encapsulins can not only encapsulate endogenous but also exogenous cargo, thanks to an extension sequence in C-terminus.^14,16^ The cumulation of all these properties makes encapsulins promising candidates for developing drug delivery carriers, imaging and diagnosis agents or bio-nanoreactors. In our studies we focus on the encapsulins of two species: *Thermotoga maritima* (Tm) and *Brevibacterium linens* (Bl). The goal here is to perform an in-depth investigation to further extend their surface modification possibilities in order to introduce functionality and improve stability for Tm encapsulin (Tmenc) and Bl encapsulin (Blenc). Additionally, we aim to increase the purification yield of encapsulin for use at larger scale.

Encapsulins are protein cages composed of 60 monomers assembled in a T=1 symmetry (a cage is composed of 60 asymmetric units made of one protein).^14,17^ The encapsulins’ surface can be modified either chemically or genetically. The latter allows insertion of a desired peptide (e.g., a targeting peptide) with high control over both the amount and the position on the cage. This peptide insertion at a single site of the protein monomer will lead to 60 functional groups on the nanocage surface. The design and fine tuning of modifiable locations, without disturbing the cage structure, allows for versatile surface modifications, which is crucial for a large variety of applications.

So far, seven genetically modified sites have been reported on encapsulins, which were all developed on the Tmenc.^18–22^ The monomeric protein of Tmenc consists of three domains; P, E and A (Figure 1B). The reported modifications are located on the N-terminus and in loop positions 42-43 (which are in the P-domain), 138-139 and C-terminus (A-domain), and loop positions 57-58, 60-61, 71-72 (E-domain). Moreover, among those seven positions five are exposed on the surface of Tm encapsulin, which are located in the A and E-domain.^18,22^ For these five surface exposed positions, results from Moon et al. suggested that the C-terminus is less accessible. The variant with 6 histidines inserted at the C-terminus could not be purified on nickel column.^18^ Regarding modifications in the E-domain, the modification between the residues 71-72 leads to a disruption of the cage, while modifications after residues 57 and 60 are mostly insoluble with low purification yield.^22^ Clearly, there is a need for further investigations about alternative loops that may be engineered on the surface of Tmenc, and by homology on Blenc. In this study, we reveal two new positions on the surface of Tmenc where a peptide can be inserted and we extend the C-terminus with a peptide of an additional 10 amino acids. Furthermore, an improved protocol for encapsulin purification is described. Noticeably, our study demonstrates that the sequence homology between different encapsulin species doesn’t exactly reflect the structural homology and we show the possibility of using the C-terminus as an engineerable position on the surface of Blenc.

**Figure 1.**
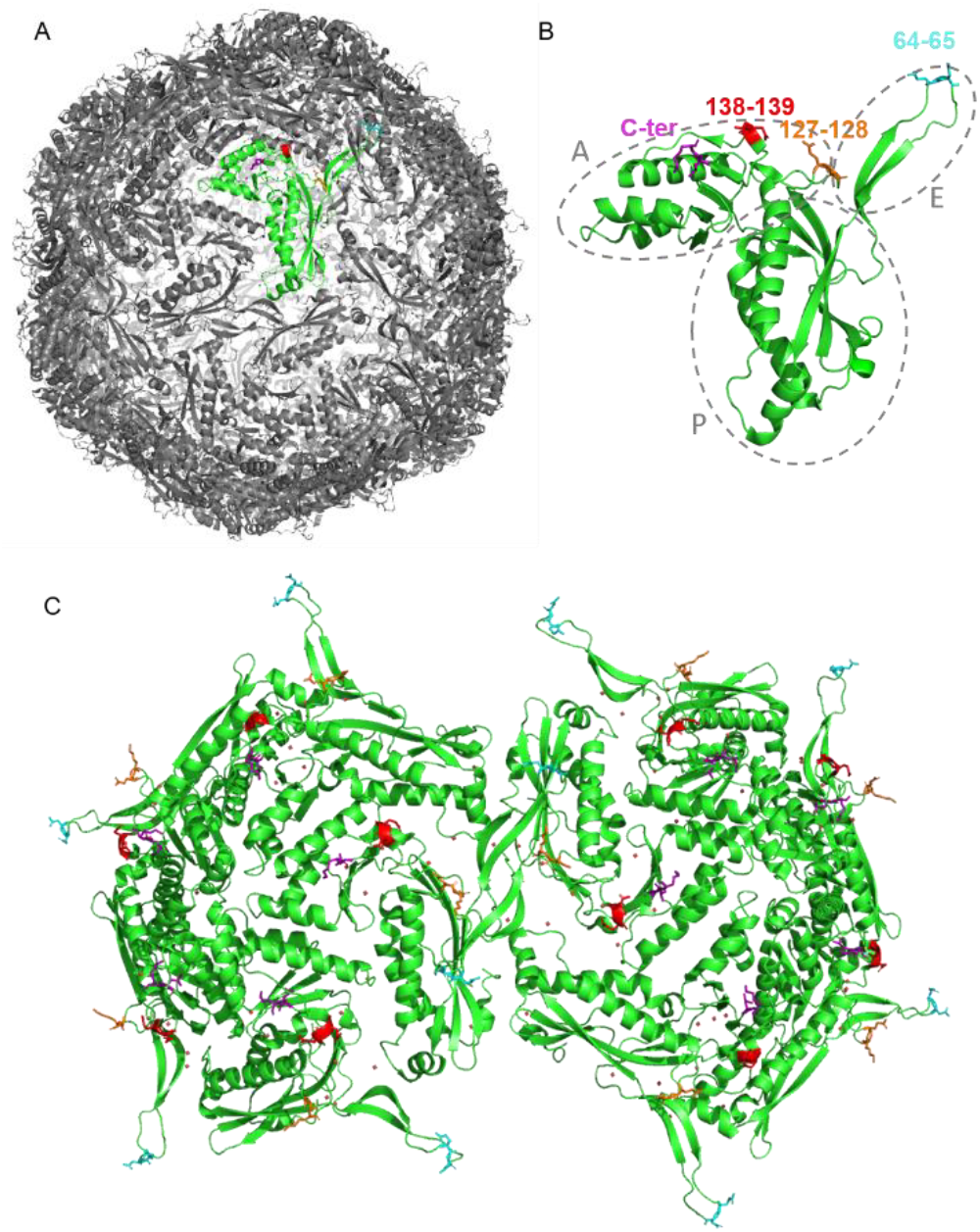
Structure of Tm encapsulin (Sutter et al.^13^) highlighting loop 64 (cyan) with glutamate 64 and asparagine 65. Loop 127 (orange) with glutamate 127 and lysine 128. Loop 138 (red) with glutamate 138 and 139. The C-terminus is in purple with the two terminal residues being lysine and phenylalanine. **A** The entire cage is shown with one monomer in green. **B** One monomer with the positions of the different exposed loops. **C** Zoom in of two pentamers, to show the different positions of the loops on the surface of the encapsulin.

## MATERIALS AND METHODS

### Bacterial strains & Plasmids

Bacterial strains and plasmids used in this study are listed in Table 1.

**Table 1.**
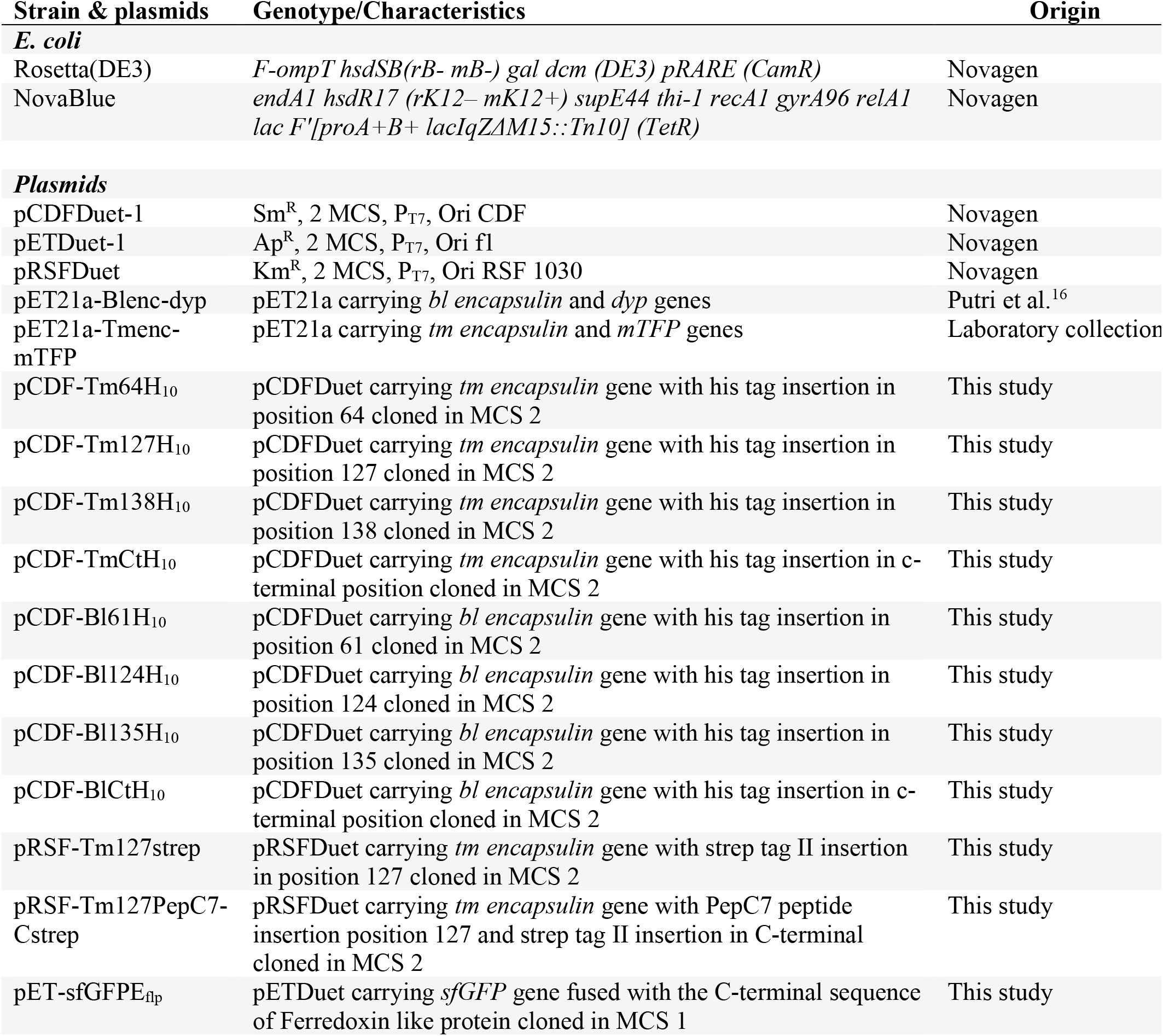
Bacterial strain and plasmids used in this study

### DNA manipulation

Plasmid preparation, DNA purification and PCR product purification were performed using the appropriate Macherey Nagel kits. Restriction enzymes, DNA polymerase and other molecular biology reagents were purchased from New England Biolabs. High fidelity polymerase Q5 (NEB) was used for PCR amplification. The sequences of all used oligonucleotides (purchased from EurofinsGenomics) are listed in Table 2. To construct the expression plasmids for encapsulin gene, the encapsulin genes were PCR-amplified using corresponding primers, and cloned into pCDFDuet or pRSFDuet vectors (Novagen) using SLIC method^23^ between NdeI-EcoRV (MCS2) restriction sites. The sfGFP gene was cloned into pETDuet vector (Novagen) using SLIC method between NcoI-SalI (MCS1). The sequences of all plasmids were verified using the sequencing service of EurofinsGenomics.

**Table 2.**
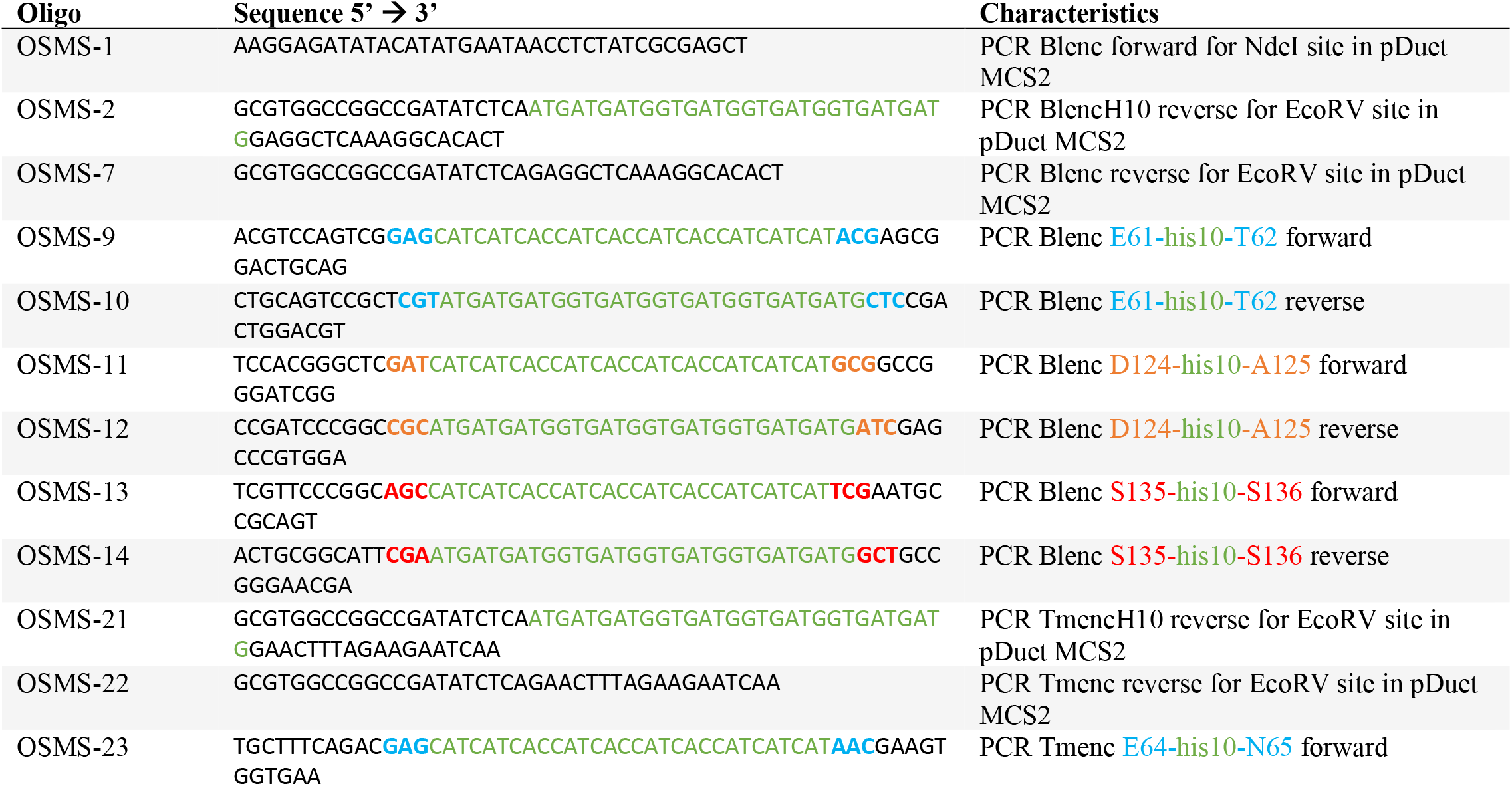
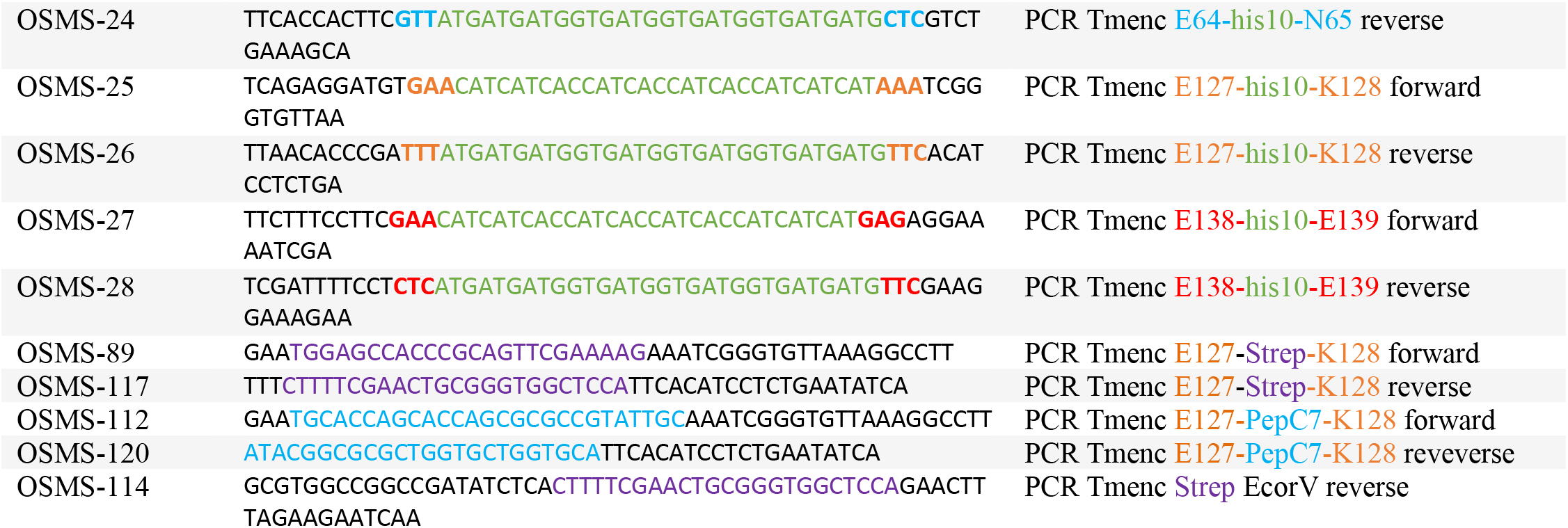
Oligonucleotides used in this study. Dna sequence is colored corresponding to the residue or peptide sequence described in characteristics

### Protein production and purification

Competent cells of *E. coli* Rosetta strain were transformed with pCDF or pRSF-enc variants or co-transformed with pRSF-enc variants and pET-sfGFPE_flp_. The bacteria were grown until 0.5 OD at λ = 600 nm at 37°C in LB medium with appropriate antibiotic (Streptomycin 30 μg/ml, Kanamycin 30 μg/ml and/or Ampicillin 50 μg/ml). The expression of the different encapsulin genes was induced with 1 mM of IPTG for 12 h at 25°C. Bacteria were collected by centrifugation and lysed by sonication (2 × 1 min) in 50 mM Hepes buffer (pH 8, 150 mM NaCl, 1 mM EDTA, 20 mM MgCl_2_, 1 protease inhibitor tablet/7mL (cOmplete^TM^), 0.5 mg/mL Lysosyme, 20 μg/ml DNAse, 30 mM Imidazole, 15 mM Beta-mercaptoethanol (βme)). The lysate was cleared by ultracentrifugation (20000 x g) to remove cell debris. Cleared lysates containing encapsulin histidine variants were each loaded onto separate 3-ml Ni-NTA-functionalized agarose beads (Protino® Ni-NTA Agarose) in Biorad gravity columns for 1 h. The immobilized proteins were washed using washing buffer (50 mM Hepes pH 8, 150 mM NaCl, 15 mM βme and 30 mM imidazole) and eluted in elution buffer (50 mM Hepes pH 8, 150 mM NaCl, 15 mM βme, 500 mM imidazole). The buffers for the strep encapsulin variant purification were not containing imidazole. The cleared lysate containing encapsulin strep variants were loaded on StrepTrap Hp 5 mL column (Cytiva) using a BioRad NGC FPLC. The immobilized proteins were eluted in elution buffer (50 mM Hepes pH 8, 150 mM NaCl, 15 mM βme, 2,5mM desthibiotine).

The proteins were concentrated using Amicon^®^ Ultra Centrifugal filters (Millipore, 100-kDa cut-off) and purified using size exclusion chromatography (SEC) on a Superose 6 10/300 increase column pre-equilibrated with 50 mM Hepes pH 8, containing 150 mM NaCl and 15 mM βme.

### SDS-PAGE and immuno-detection

Proteins from different purification steps were separated by electrophoresis on 15% polyacrylamide gels and stained using Coomassie brilliant blue. Proteins from bacterial extracts were separated by electrophoresis on 15% polyacrylamide gels and transferred onto nitrocellulose membranes using a wet blotting apparatus (BioRad). Membranes were blocked with 5% milk in PBST (Phosphate buffer saline, 0.05% Tween 20) and incubated with monoclonal mouse anti-His antibody (Penta His, Qiagen, dilution 1:1000) according to the manufacturers’ instructions. This was followed by two 10 min washes and a 1 h incubation in peroxidase-labelled anti-mouse antibody (1:2000, Sigma). Membranes were developed by homemade enhanced chemiluminescence and scanned using FluorChem M hardware (Proteinsimple).

### Transmission electron microscopy (TEM)

TEM measurements were performed on a Philips CM300ST-FEG Transmission Electron Microscope. 5 μL of each sample was applied to a Formvar carbon-coated copper grid (Electron Microscopy Sciences). Samples were incubated on the grid for 2 min, then any excess buffer was removed with filter paper. Samples were negatively stained by applying 5 μL uranyl acetate (1% w/v) onto the grid and incubating for 40 s. Any excess stain was removed, and the samples were left to dry for 10 min before imaging. The size of encapsulin cages was determined by using ImageJ (http://imagej.nih.gov/ij/) and making the mean on 30 measurements.

### Dynamic light scattering (DLS)

The hydrodynamic size distribution of the particles was determined using a Nanotrac Wave (Microtrac) particle analyzer. An average over 5 runs of 120 s each was used to determine the size deduced from the intensity distribution. The number distribution was used to visualized where is the major contribution.

### Cellular uptake

bEND.3 cells (ATCC) were seeded at 10×10^3^ cells per well in a 96-well plate in Dulbecco modified Eagles medium (DMEM), fetal bovine serum (FBS), penicillin-streptomycin (containing 10.000 units penicillin, 10 mg streptomycin mL-1). After 24h incubation at 37 °C in humidified 5% CO2-containing atmosphere TmPepc7-sfGFP and TmStrep-sfGFP were diluted to 50 nM in DMEM and added to the cells. The particles were incubated for 4h. Subsequently, the cells were washed with phosphate buffered saline (PBS, pH 7.4) with 4% Paraformaldehyde (PFA) and stained with 4,6-diamidino-2-phenylindole dihydrochloride (DAPI) and Wheat Germ Agglutinin (WGA) staining. The cellular uptake was analyzed by fluorescent microscopy on an Olympus IX2-ILL 100 with 20x objective. The filters used were ex 350/em 420 for DAPI/WGA and ex 460-490/em 525 for GFP.

## RESULTS AND DISCUSSION

### Selection of loops on Tmenc and Blenc and variant construction

To construct encapsulin variant we first select promising loops candidates for surface modifications. This selection can be done using the resolved encapsulin structure. For Tmenc the structure has been described by Sutter and collaborators^14^ and we used the PyMol software (The PyMOL Molecular Graphics System, Version 2.1 Schrödinger, LLC). Following this approach, loops at positions 64, 127, 138 and the C-terminus of Tmenc were investigated (Fig. 1). As modifications on the loop at position 138 and C-terminus were already characterized by Moon and collaborators,^18^ we decided to use position 138 as a benchmark for our experiments to check that we could reproduce their result. In addition, while Moon *et al* couldn’t purify the variant when modifying the C-terminus, we investigated the possibility of getting encapsulin purification using a longer 10 histidine tag instead of 6. Indeed, increasing the length of the tag might make it accessible from the surface. The complete cage of Tmenc is represented in Figure 1A with a single monomer highlighted in green. In Figure 1B, a monomer with the designed modification loops is shown. Additionally, using a representation of a double pentamer, Figure 1C highlights the exposed position of the chosen residues. As the atomic structure of Blenc is unfortunately unavailable so far, we proceeded by homology with Tmenc. Following an alignment of sequence procedure using the ENDscript server,^25^ we selected the corresponding residues of Blenc, which are residues 61, 124, 135 and the C-terminus (Figure 2). Next, we genetically inserted 10 histidines between each set of highlighted amino acids.

**Figure 2.**
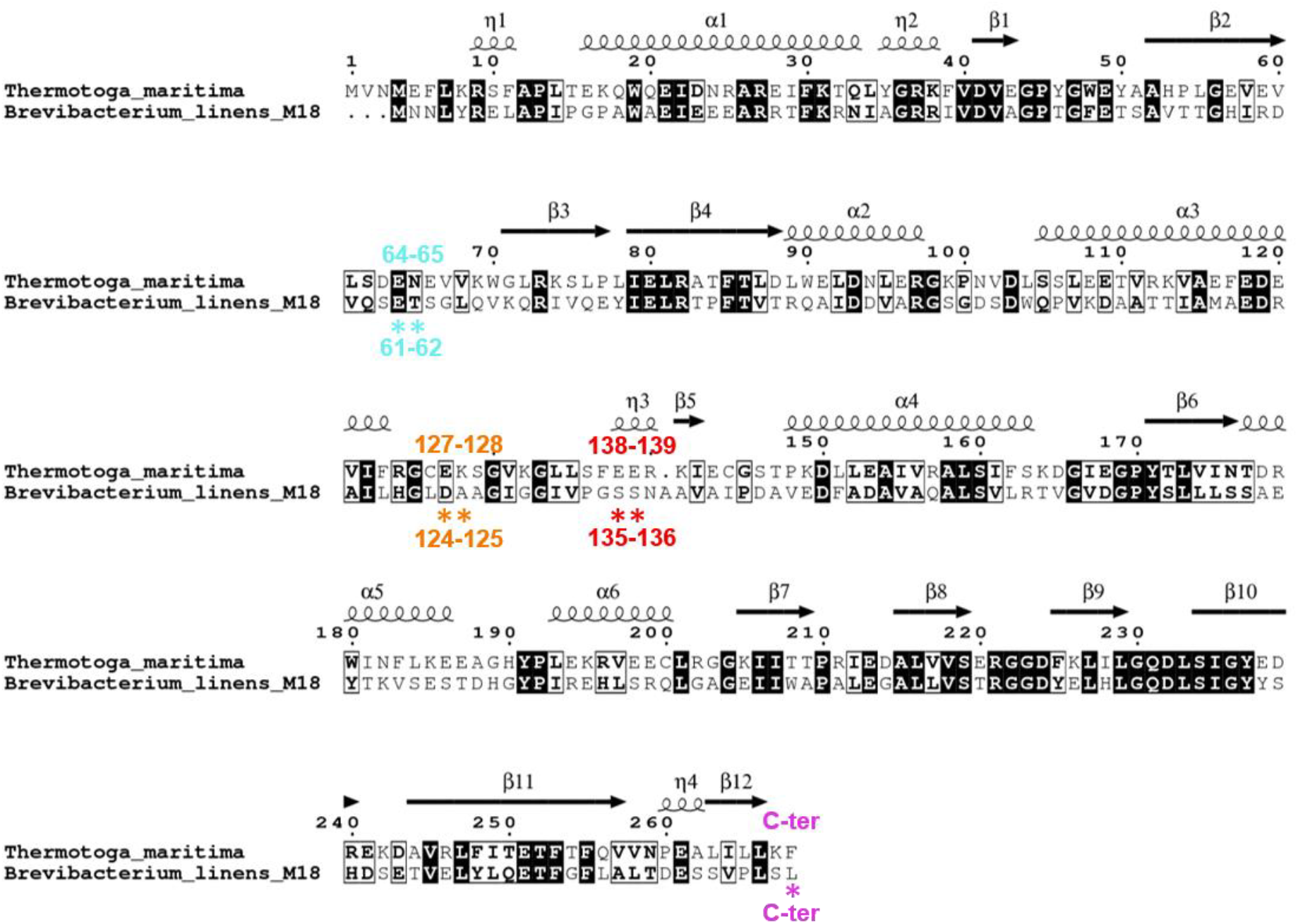
Alignment of Tm and Bl encapsulins with in black the similar residues and in boxes the homologous substitution. The stars represent the similar residues between Blenc and Tmenc for the selection of the loop on Blenc.

### Purification of the encapsulins

The variants were produced and assembled in bacteria. The soluble fraction was extracted and purified by affinity chromatography on Nickel-NTA columns. All the variants of Tmenc were purified and found in the elution fraction with a molecular weight of 32,2kDa (Fig. 3A), which implies that the new variants are stable and the His-tag is exposed on the surface. Note that multimers of Tmenc were observed on SDS-PAGE, implying that a part of Tmenc resists denaturation of the sample during preparation for SDS-PAGE analysis (heating and SDS treatment). Concerning Blenc, while BlencH (Blenc with 10 histidine in C-terminus) is correctly purified and found in the elution fractions with a molecular weight of 30kDa (Fig. 3B top), this is not the case for the other variants of Blenc. Blenc61H is purified but at a lower yield compare to BlencH, and for the variants Blenc124H and Blenc135H, no proteins are present in the elution fractions. To clarify what happens with the latter variants and why they could not be purified, we tested both their production and stability in bacteria cells. Thereby, we followed the production of Blenc124H, Blenc135H and BlencH as positive control (Fig. S1). This revealed that all the variants are produced, implying that there is no problem with the protein production. Although the stability of variants 124 and 135 differ compare to variant BlencH, proteins are still present 21 hours after production. Note that the non-attachment to the nickel column could result from either protein aggregation, forming inclusion bodies, or from the tag being hidden. The latter would suggest that the chosen loops are not exposed on the surface, or alternatively that the insertion of the His-tag involves a conformational change which buries the loop with the tag.

**Figure 3.**
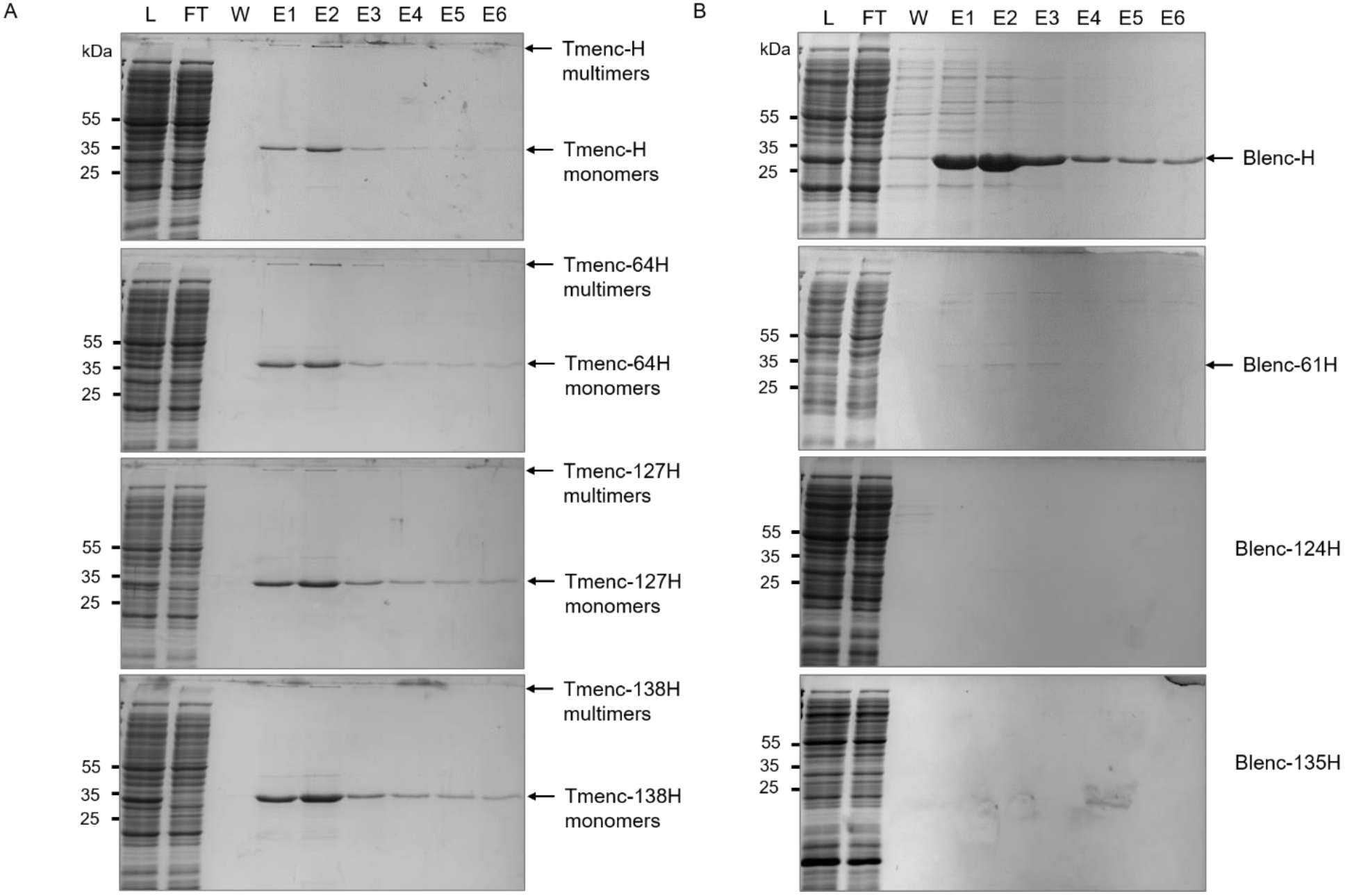
SDS-PAGE analysis of the purification of the different encapsulin variants by affinity chromatography on nickel columns. The fractions are analyzed by SDS-PAGE and stained by Coomassie blue. L = Loading, FT = Flow Through, W = Wash, E = Elution. Molecular weight markers (in kDa) are indicated on the left. The different variants are indicated on the right of the arrows except for Blenc124H and Blenc135H (B bottom), where no protein is visible. **A** Analysis of Tmenc variants. **B** Analysis of Blenc variants.

### Structure and size characterization of the encapsulins

After obtaining all the variants of Tmenc and BlencH we did a second purification step by size exclusion chromatography (Fig. S2). Note that, as previously mentioned, the other Blenc variants couldn’t be isolated and that for Blenc61H the yield was too low to performed this next step of purification. This purification step also yields additional information on the approximate size of the encapsulins. With a sepharose 6 10/300 column used in previous protocol as last purification step, it has been shown that native encapsulin elutes at V ~ 12 mL,^17^ furthermore, we did a calibration of the column and V = 12 mL correspond to a size of approximatively 2000 kDa being close to the calculated encapsulin mass (1932 kDa). To further analyze the size and morphology of the designed encapsulins we used DLS and TEM (Fig. 4). DLS measurement show a peak for a cage diameter of 25,5 nm (mode value of the distribution). From TEM image analysis the mean diameter is found to be around 21,5 nm. This difference can be explained considering the following: for TEM sample the grid preparation could lead to a “drying effect” leading to smaller particles compare to when they are in solution; for DLS the bin width is big (4,5nm) and make the size determination less precise. Most importantly, all the studied variants have a size close to the one structurally determined by Sutter *et al* and have a spherical shape similar to the wild type encapsulin observed in previous studies.^14,17,18^ Note that from the DLS data (Figure 4 D-E), we can see that the particles can cluster to form aggregate of much larger size (200-400nm). This effect is however negligeable, as revealed by the number distribution issued from DLS measurements showing that the majority of the population is in the 25 nm peak (Figure S3).

**Figure 4.**
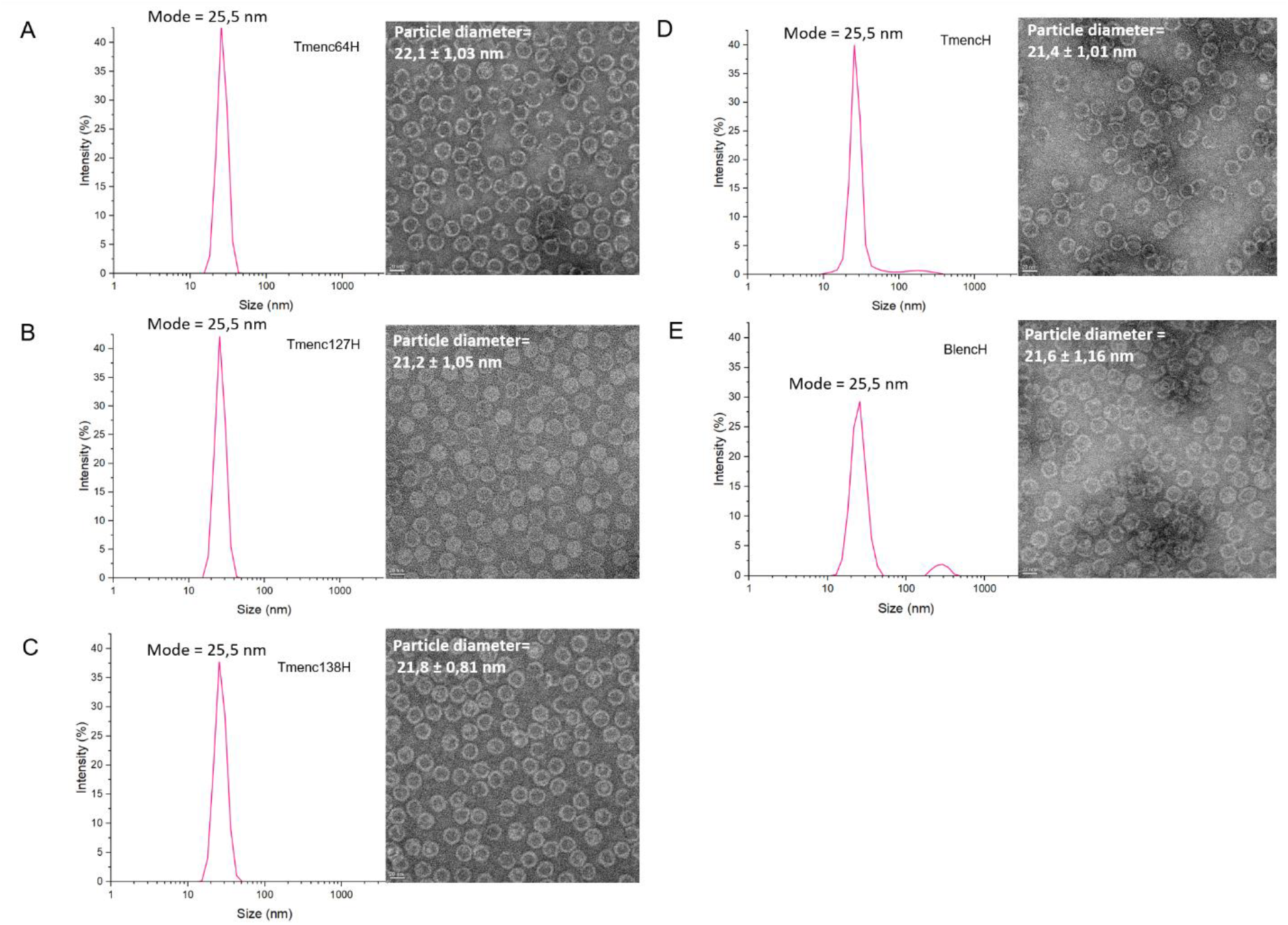
Size and morphological characterization of the Tmenc variants and BlencH. DLS-based size distributions (left) and negatively stained TEM images (right) of encapsulin particles for Tmenc64H (**A**), Tmenc127H (**B**), Tmenc138H (**C**), TmencH (**D**), and BlencH (**E**).

### Buffer optimization to increase encapsulin stability and avoid protein precipitation

Buffers used in previous studies such as Phosphate buffers, ^18,20^ Tris buffers with NH_4_Cl-MgCl_2_ or NaCl complemented with 1 mM of β-mercaptoethanol,^16^ resulted in precipitation of proteins after 1 to 3 days or during handling (Fig. S4). By changing the buffer composition to 50 mM Hepes buffer, 150 mM NaCl with 15 mM β-mercaptoethanol, we obtained stable particle solutions of all variants when stored at 4°C. Every variant solution was stable for at least 5 days and could be recovered after Amicon^®^ concentration (Fig. 4). The size of the cages was checked by DLS after 1 year of storage yielding different stabilities depending on the variant (Fig. 5). The variants Tmenc127H, Tmenc138H and BlencH are stable after 1 year of storage, while Tmenc64H showed partial aggregation. The encapsulin with the modification on C-terminal (TmencH) completely aggregated over this time frame. The number distribution shows for all variants except TmencH that the majority of the population is still forming a cage of 25 nm (Figure S5). Note that as the same His_10_-tag was inserted in all designed encapsulins, it implies that the difference in stability originates from the position of the modification and not from the nature of the modification itself. Furthermore we noticed that using our protocol we get a higher purification yield compared to the study of Lee and collaborators.^22^ From 1 L of culture, we obtained between 1 and 2 mg/mL of purified protein, while Lee and collaborators reported at best a yield of 0,039 mg/mL from 0,8 L of culture. This means a remarkable net increase of cage purification rate of about 20 times. This phenomenon could be explained by the increase of stability. However, we changed various parameters such as the buffer, the vector and the purification method compared to the work by Lee and collaborators. An in-depth investigation of the parameters which lead to such noticeable increase of the cage production rate is beyond the scope of the present work but will be addressed in a further study.

**Figure 5.**
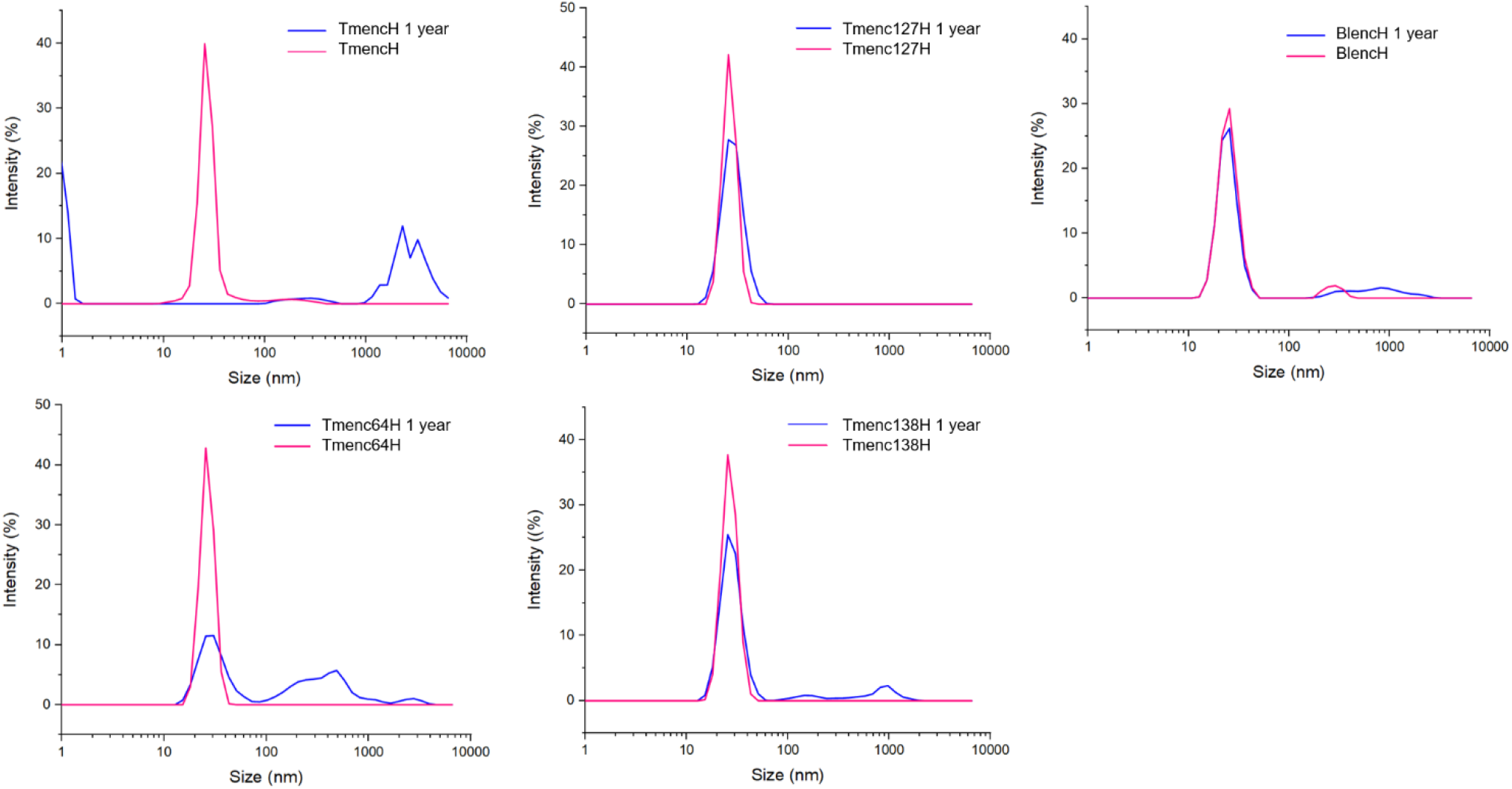
Stability study of different encapsulins variants by DLS. Size distributions of Tmenc64H, Tmenc127H, Tmenc138H, TmencH, and BlencH after 5 days (pink line) and 1 year (blue line).

### Functionalization with PepC7 targeting peptide

To demonstrate the use of multi-functionalized encapsulin for different applications, the targeting peptide PepC7 was added in position 127 of Tmenc. In addition, to keep the advantage of affinity chromatography purification, a Strep-tag II was added in C-terminus. This peptide PepC7 (CTSTSAPYC) was identified by phage display as a brain targeting peptide.^26,27^ The brain drug delivery is a big challenge, due to the difficulty of getting through the Blood Brain Barrier (BBB). One of the emergent strategy is the BBB shuttle peptide which can be combined with nanoscale drug delivery carrier.^28–30^

To follow the encapsulin uptake by the brain cells a fluorescent protein, the super-folder GFP (sfGFP), was fused with the E extension (C-terminus sequence) of the native cargo, Ferredoxin like protein (Flp), to be encapsulated in the cage.^14^ The encapsulin Tm127PepC7/sfGFP was purified by affinity on Strep-Tactin column following by a size exclusion chromatography similarly to the purification process of the Tmenc histidine variants (Figure S6 B). This is an important result as it demonstrates that position 127 and C-terminus can be both modified simultaneously.

The uptake was tested on brain epithelial cells (bEnd.3 cells). As a control of PepC7 efficiency, a cage with Strep Tag modification only was engineered, loaded with the sfGFP and purified (Figure S6 A). One of the advantage of using Tmenc for targeting drug delivery system is that Tmenc it is almost not uptaken if it does not have specific surface modification, as already observed in previous studies with different cell lines.^31^ Consistently, similar behavior is observed here with the bEnd.3 cell line. There is almost no uptake of TmStrep/sfGFP and we observe an uptake (green spots) of TmPepC7/sfGFP by the cells (Figure 6). This demonstrates the possibility of targeting the BBB with encapsulin modified with PepC7 targeting peptide making it a useful tool for further investigation on this topic.

**Figure 6:**
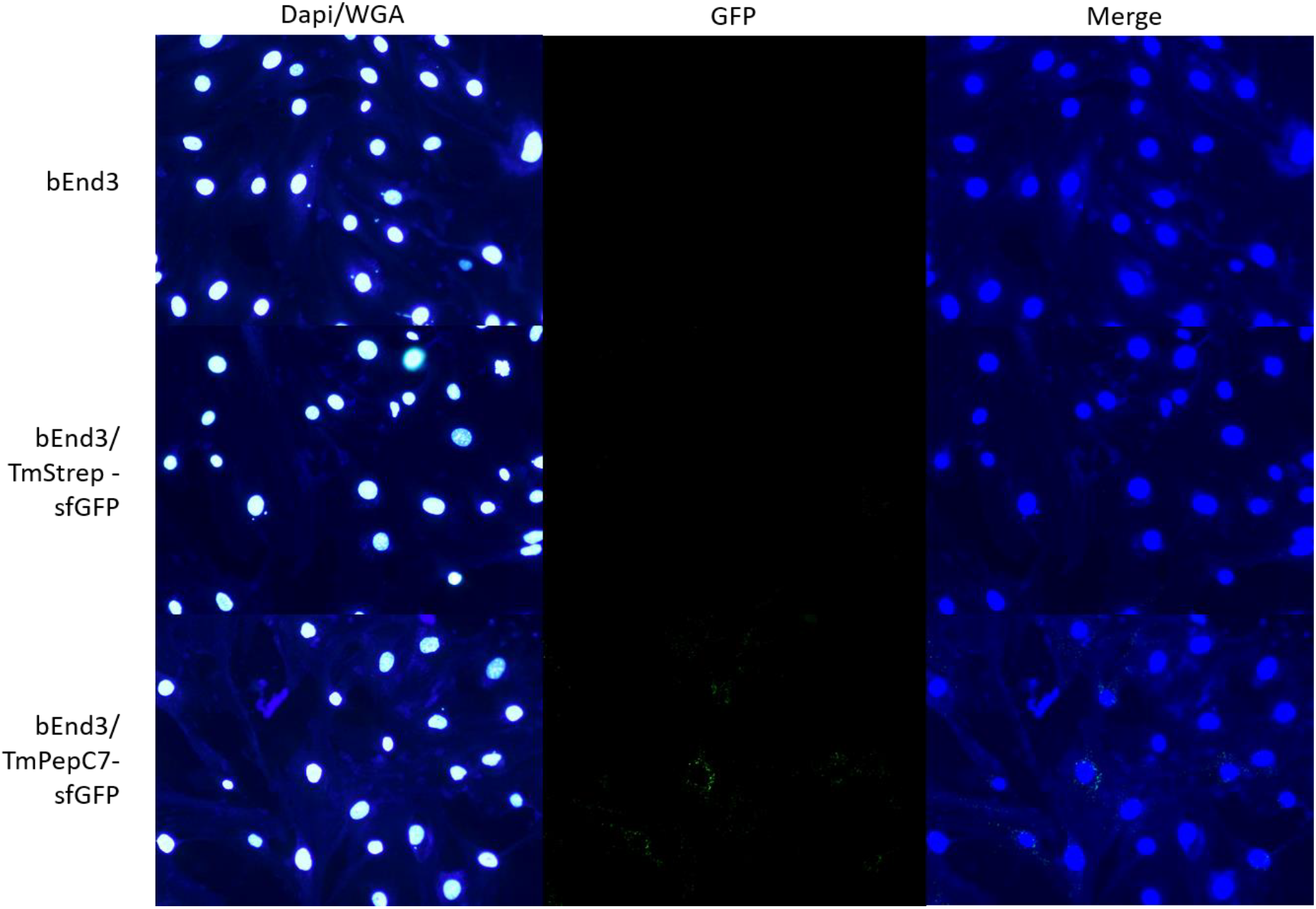
Tmenc uptake by bEnd.3 cells. Fluorescent microscopy images with in first row the cell untreated, the second raw the cells treated with 50nM of Tm127strep/sfGFP and the third raw the cells treated with 50nM of Tm127PepC7-Cstrep/sfGFP. The first column shows the DAPI (nuclei) and WGA (membranes) staining (filter 350-420), the second column show the GFP emission (filter 460/90-525) and the third the merge of these two channels.

## CONCLUSION

This study reports the successful insertion of functional loops at 2 new positions on the surface of Tm encapsulin, that is after residue 64 and 127. It also shows the successful modification of the C-terminus of Bl encapsulin which is exposed and accessible on the surface. Modification of the C-terminus of Tmenc was also investigated, and we show that compared to the study of Moon and collaborators,^18^ an increase of the size of the histidine tag from 6 to 10 residues allows the purification by immobilized metal affinity chromatography. Consequently, this position can be used for further applications by adding a linker between the inserted peptide and the encapsulin.

As the structure of Blenc is not available, we used a sequence and structure homology approach between Tmenc and Blenc to investigate modifiable loops on Blenc. However, Blenc variants could not be isolated, implying that we cannot engineer encapsulin cages using an homology approach with Tmenc: the structure is essential for an efficient investigation of new modifiable positions.

Using our protocol, we were able to considerably increase the short-term stability of encapsulins and we showed that some sample can have long-term stability up to a year. This study demonstrates that even if the integrity of the cage is kept, some modifications on the surface affect the structure and induce differences in long-term stability.

The utilization of encapsulins for medical applications is highly promising thanks to their large set of advantages resulting from their small size and their biocompatibility. Note that further studies are yet required to fully evaluate the encapsulin immune reaction. Previous studies already showed how encapsulin can be employed for liver targeting, vaccine development, imaging or as nanoreactors.^6,20,32–34^ Therefore, it is crucial to deepen our knowledge about encapsulin modification and to optimize their production and usability. Hence, this study paves the way for the development of improved encapsulin engineering and production. It should enable a wide range of further studies investigating multivalency, targeting and delivery for multi-vaccine or multi-targeting applications. Indeed, further studies are ongoing to create well-defined modified surfaces by having 60 functional groups when every protein subunit has a single modification and 240 functional groups when each subunit has 4 (different) modifications. So far in this study we demonstrate the possibility to have two positions modified simultaneously, the position 127 and the C-terminus leading to a protein nanocage with 180 functional groups.

Finally, we demonstrated the possibility of targeting the BBB with encapsulin. Future investigations using different BBB shuttle peptides and BBB transport model are ongoing to investigate the utilization of encapsulin for BBB drug delivery.

## SUPPORTING INFORMATION

Figure S1. Production and stability of the variants BLenc124H and Blenc135H in bacteria.

Figure S2. Purification of the different encapsulin variant by size exclusion chromatography.

Figure S3. DLS measurement of encapsulin variants using number distribution

Figure S4. Precipitation of Tmenc64H in a dialysis bag.

Figure S5. Encapsulin stability analysis using DLS measurement with number distribution.

Figure S6. Purification of Tm127strep/sfGFP and Tm127PepC7/sfGFP by size exclusion chromatography.

## ACKNOWLEDGEMENTS

We thank Dr. E. G. Keim, (MESA+ Institute for Nanotechnology, University of Twente) for assistance with TEM measurement. We thank Dr. Mathieu Souzy, Dr. Dorothee Wasserberg and Jessica Nettofrez for proof reading the manuscript.

## SUPPORTING INFORMATION

**Figure S1.**
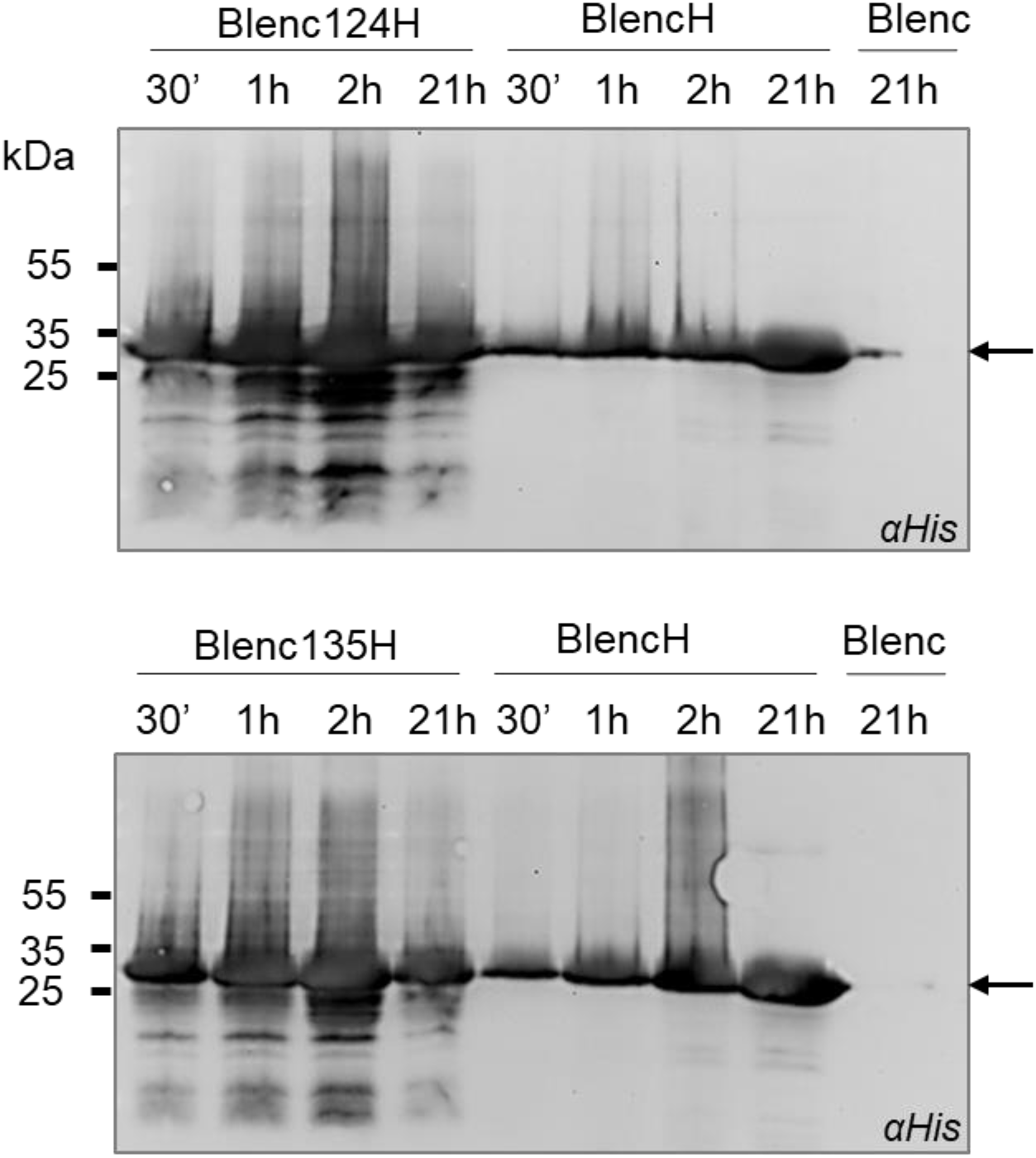
Production and stability of the variants BLenc124H and Blenc135H in bacteria. SDS-PAGE revealed by immunodetection with an alpha-His antibody directed against the His-tag. Native Blenc is used as a negative control for the immunodetection. BlencH is used as a positive control. The production and stability are checked at different time points after induction, 30min, 1h, 2h and 21h. Black arrows shows the monomers of Blenc.

**Figure S2.**
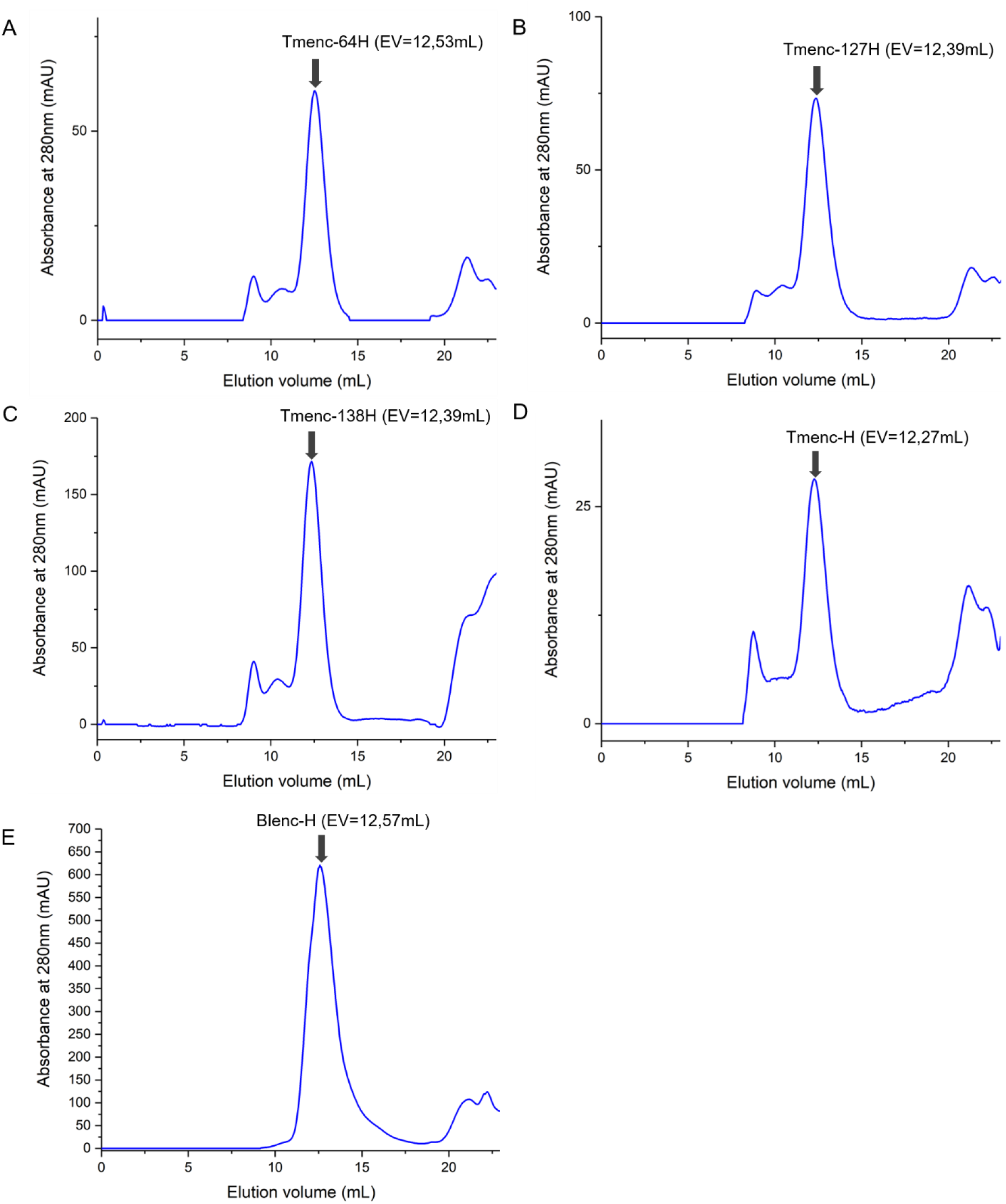
Purification of the different encapsulin variants by size exclusion chromatography. Size exclusion profile of variants Tmenc64H (**A**), Tmenc127H (**B**), Tmenc138H (**C**), TmencH (**D**), and BlencH (**E**) monitored at λ=280nm. The elution peak of the cage is shown by a black arrow around V=12mL for all variants.

**Figure S3.**
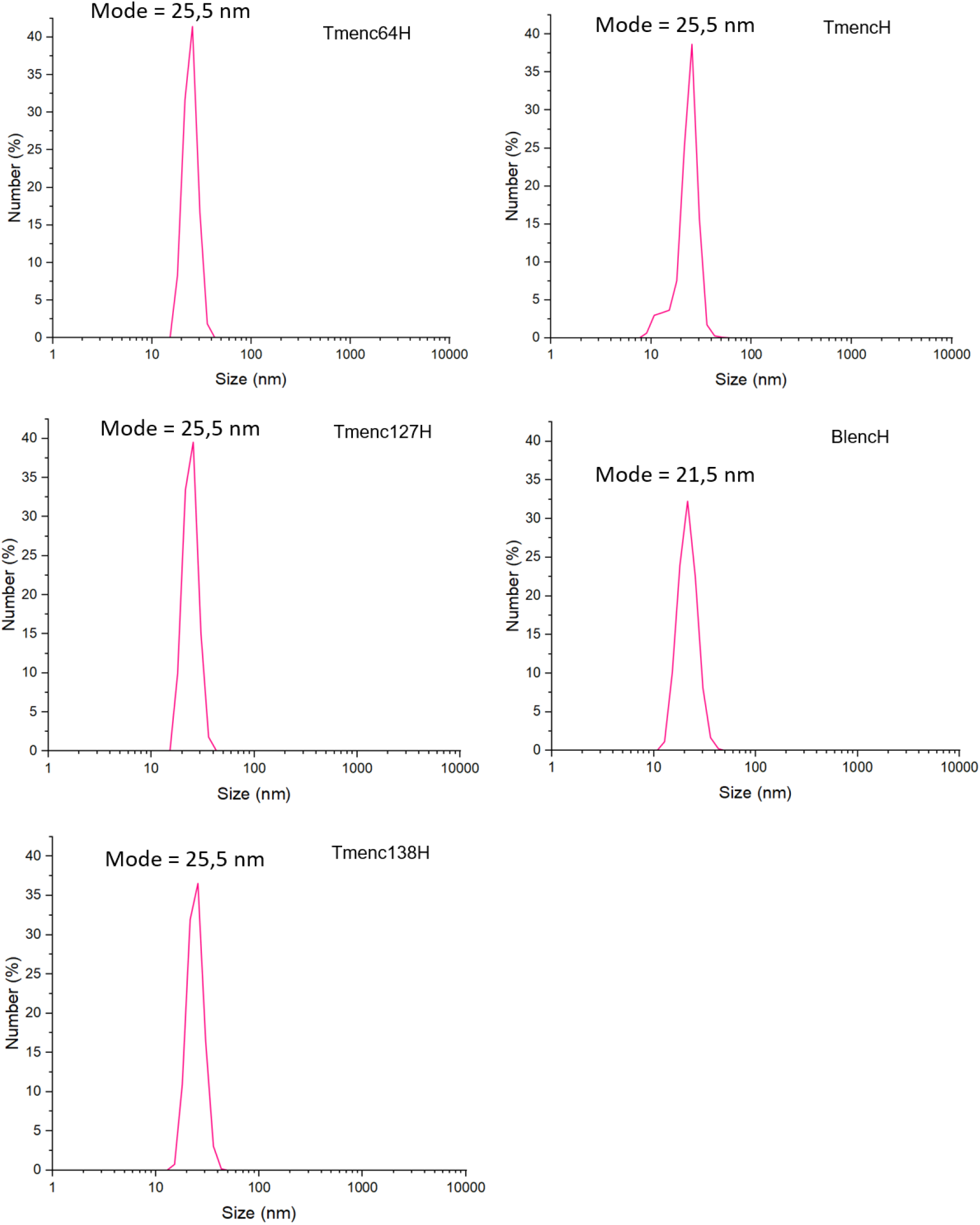
DLS measurement of encapsulin variants using number distribution. Measurments for Tmenc64H, Tmenc127H, Tmenc138H, TmencH and BlencH.

**Figure S4.**
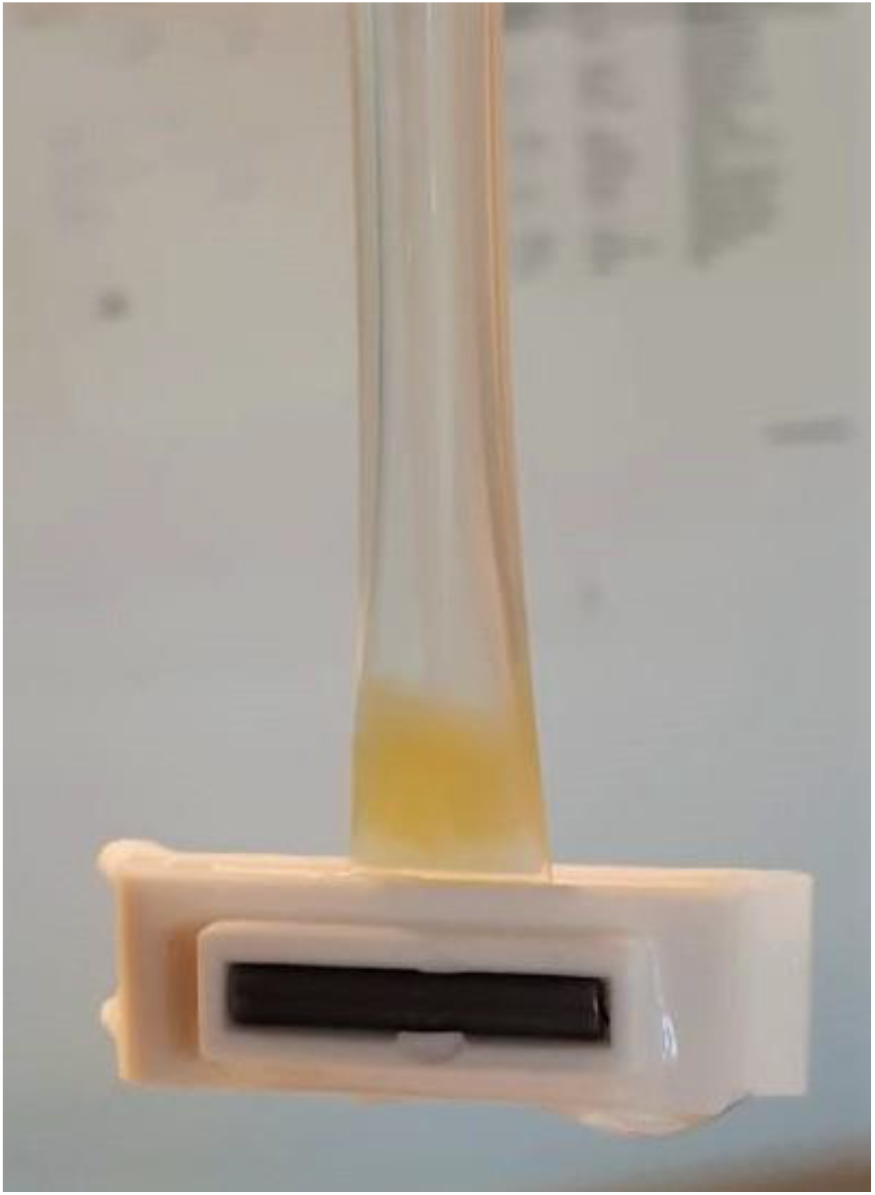
Precipitation of Tmenc64H in a dialysis bag.

**Figure S5.**
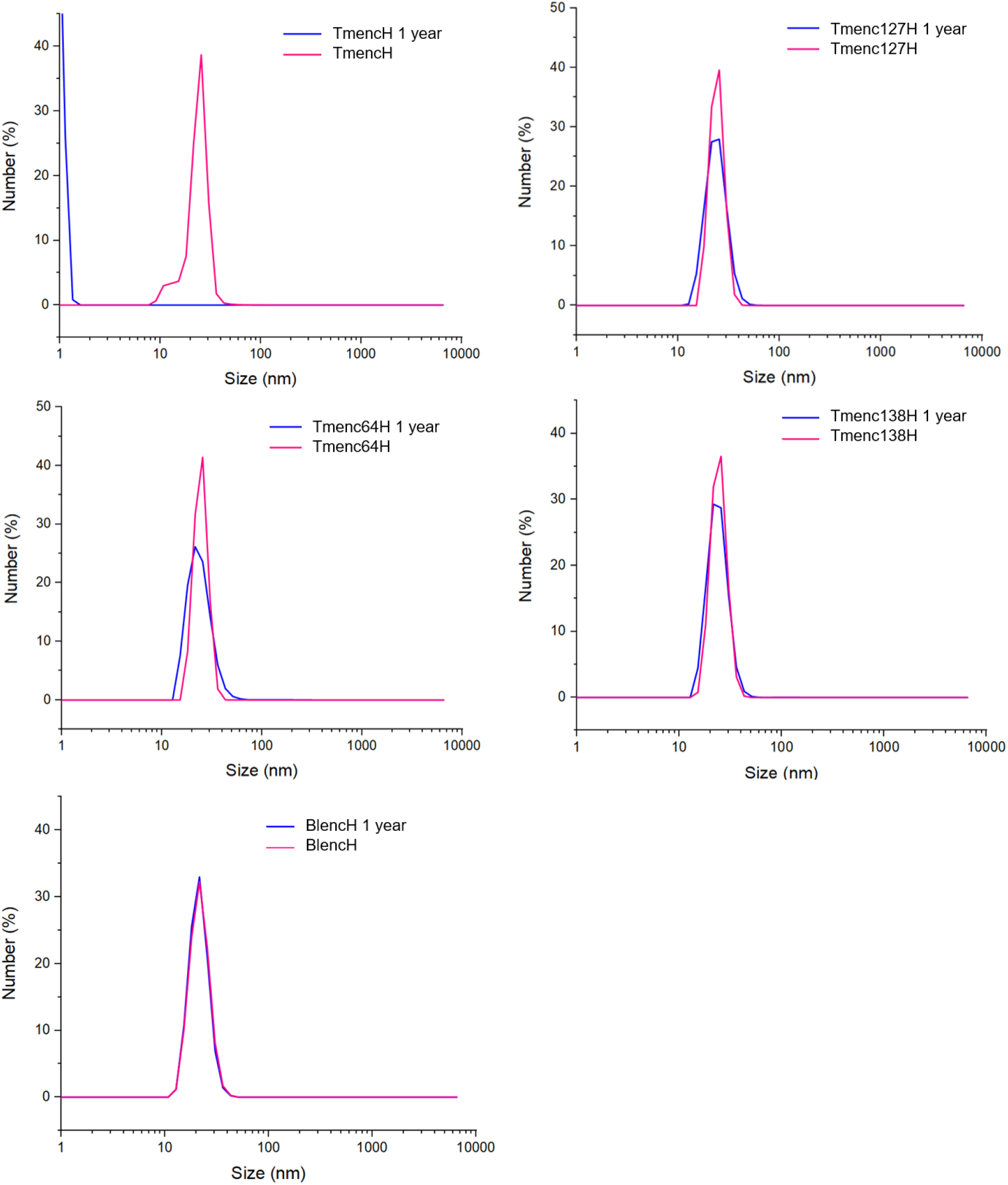
Encapsulin stability analysis using DLS measurement with number distribution. Number distributions of Tmenc64H, Tmenc127H, Tmenc138H, TmencH, and BlencH after 5 days (pink line) and 1 year (blue line).

**Figure S6.**
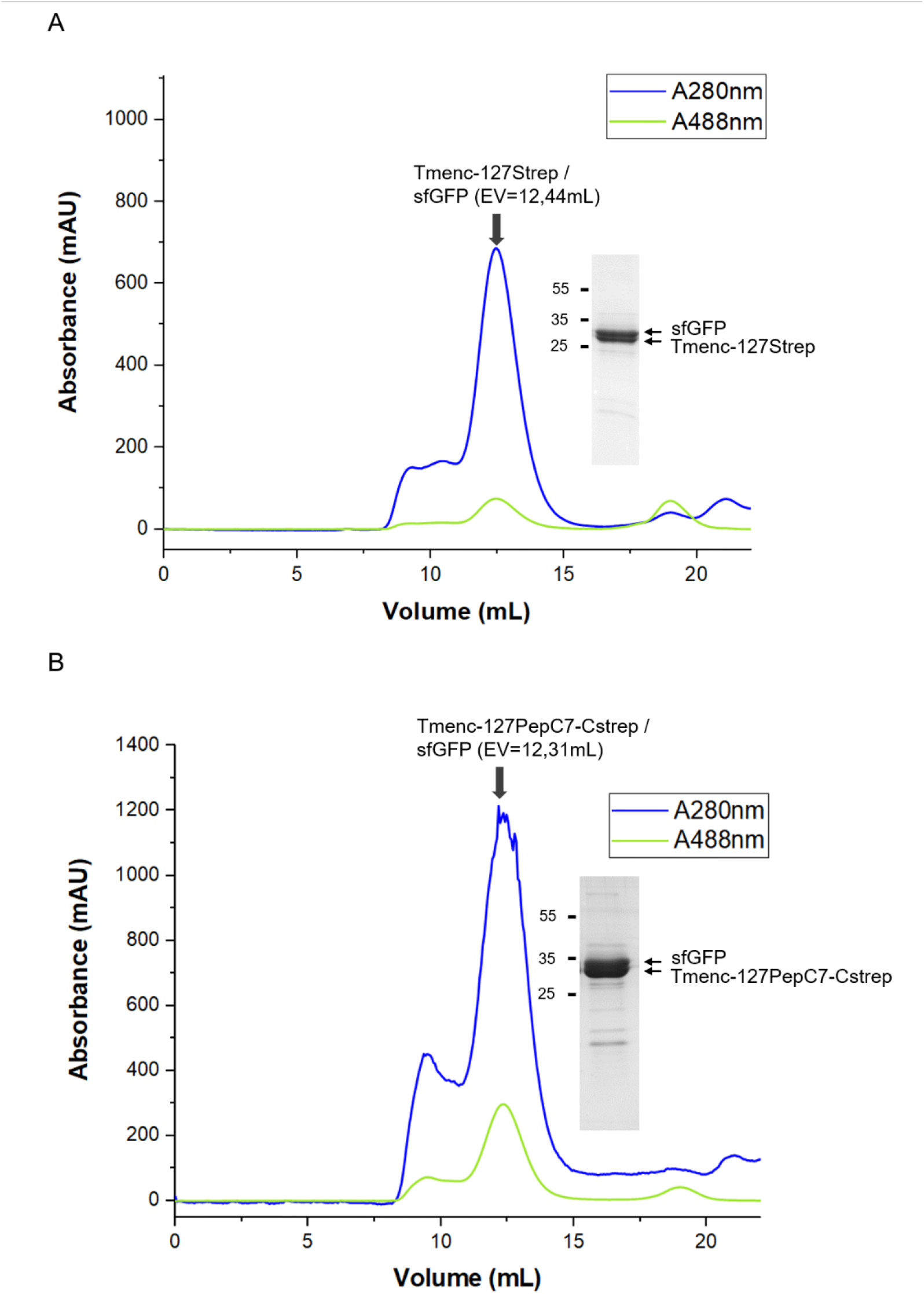
Purification of Tm127strep/sfGFP and Tm127PepC7/sfGFP by size exclusion chromatography. Size exclusion profile of variants Tm127strep/sfGFP (**A**), Tm127PepC7/sfGFP (**B**) monitored at λ=280nm for the whole proteins and λ=488nm for the sfGFP. The elution peak of the cage is shown by a black arrow around V=12mL for both variants. On the right of each peak there is its analysis on SDS-PAGE.

